# Mechanical plasticity of the ECM directs invasive branching morphogenesis in human mammary gland organoids

**DOI:** 10.1101/860015

**Authors:** B. Buchmann, L.K. Meixner, P. Fernandez, F.P. Hutterer, M.K. Raich, C.H. Scheel, A.R. Bausch

## Abstract

Although branching morphogenesis is central for organogenesis in diverse organs, the underlying self-organizing principles have yet to be identified. Here, we show that invasive branching morphogenesis in human mammary organoids relies on an intricate tension-driven feedback mechanism, which is based on the nonlinear and plastic mechanical response of the surrounding collagen network. Specifically, we demonstrate that collective motion of cells within organoid branches generates tension that is strong enough to induce a plastic reorganization of the surrounding collagen network which results in the formation of mechanically stable collagen cages. Such matrix encasing in turn directs further tension generation, branch outgrowth and plastic deformation of the matrix. The identified mechanical feedback-loop sets a framework to understand how mechanical cues direct organogenesis.

---

Branching morphogenesis leads to the formation of a tree-like, ductal epithelium in diverse organs such as kidney (1), lung (2), pancreas (3) or mammary gland (4) and is known to be the result of an interplay between chemical signals and mechanical cues (5, 6). In particular, structural and mechanical properties of the surrounding extracellular matrix seem to act as guiding cues for branch elongation (7–9). However, the mechanisms guiding the collective outgrowth of epithelial structures have yet to be identified (10–12). As branching morphogenesis differs greatly between tissues due to organ-specific mechanical and chemical environments (5, 13), the self-organization processes leading to ductal elongation remain a topic of intense investigation. Tip-cell driven invasion, as observed during *Drosophila tracheal* tube elongation (14) or retinal endothelial sprouting (15), has been described as one possible mechanism. Here, it is assumed that designated leading tip cells invade the extracellular matrix (ECM), whereas stalk cells within the duct merely push outward by intercalation processes (16, 17). In contrast, for other organs such as lung (2), kidney (1) or mammary gland (18) a non-invasive elongation process was observed. Specifically, in terminal end buds of mouse mammary organoids collective cell rearrangements were shown to drive elongation of buds without a clear mechanical interaction with the surrounding Matrigel (18). In contrast in a collagenous environment, the direction of the branching epithelium of mammary gland organoids is guided by aligned collagen fibers (8, 9), highlighting the importance of the ECM as an essential cue (19–21). Yet, the underlying self-organizing principles driving the branching morphogenesis still remain elusive. This is partly due to the fact that until now only little is known on the dynamic nature of the interaction of the epithelial layers with the surrounding ECM. Especially, most times only the purely linear elastic response of the ECM is considered, without taking into account non-linear and plastic effects (22). Using human primary single cell-derived mammary organoids we identified plastic collagen fiber alignment, collagen accumulation, tension generation and branch elongation as a dynamic collective process governed by a force-feedback loop.

## Mammary gland organoids invade the ECM by non-continuous contractions

Single primary basal human mammary epithelial cells were embedded in freely-floating 3D collagen gels as previously described (23), and expansion of the developing organoids was observed by live-confocal-microscopy over extended time periods, from hours to days (Fig. 1a). We were able to distinguish three development stages (Fig. S1a): the establishment phase (day 1 to 7), the branch elongation phase (day 7 to 11, Fig. 1b) and the alveologenesis phase (day 11 to 14). Characteristic for the establishment phase were small, rod-like cell clusters with length between 150 µm and 250 µm, that showed only rudimentary branches. During the elongation phase, these branches invaded into the ECM and developed primary and secondary side branches. Ultimately, during the alveologenesis phase, the organoids reached diameters of up to 1 mm and the branches formed rounded end buds and grew thicker allowing the formation of a lumen. This phase also coincided with polarized expression of markers for the two major lineages within the mammary gland: p63 for basal cells, expressed within the outer cell layer adjacent to the ECM, and Gata-3 for luminal cells, expressed within the cell layer adjacent to the forming lumen (Fig. 1c). This expression resembled the bilayered architecture of the human mammary gland.

**Fig. 1:**
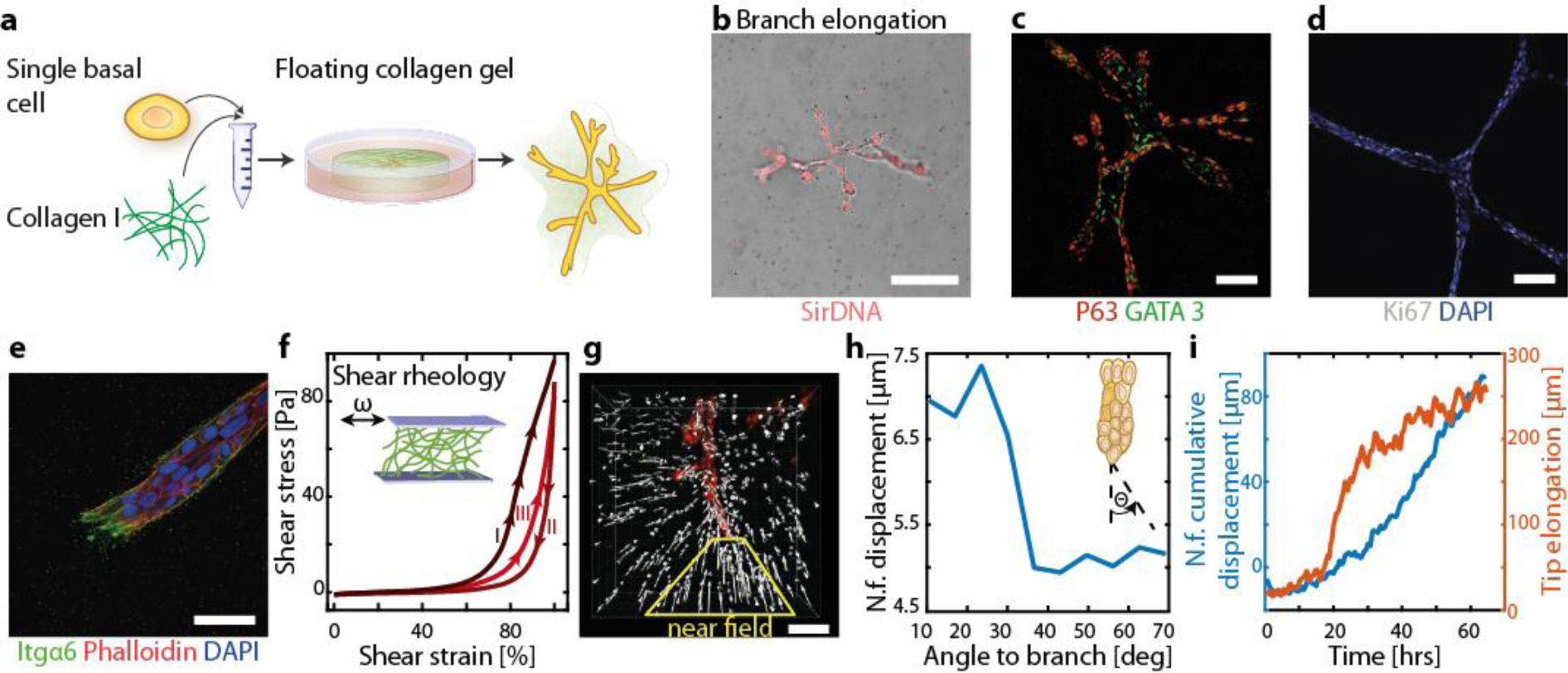
Mammary gland organoids invade the ECM by non-continuous contractions. **a** Single basal mammary epithelial cells cultured in floating collagen gels gave rise to branched organoids over a time course of 14 days. **b** Characteristic organoid morphology during the branch elongation phase including nuclei staining. **c** Immunostaining reveals a polarized expression of basal (p63) and luminal (GATA3) lineage markers at day 11 of culture. **d** Proliferative (ki67+) cells are observed throughout the whole organoid during branch elongation. **e** Leading cells invade collagen gel by actin-rich protrusions and attach via integrin-α6 to the ECM. **f** Rheology of collagen network shows Mullins-softening after cyclic strain. **g** The deformation field in the near field (n.f.) of a branch is highly anisotropic with high deformations in front of the branch which nearly vanishes at the sides of the branch. n=3 donors. **h** Representative displacement in dependency to angle of the branch. **i** The average cumulative displacement in front of a branch in the near field of a branch (blue) and the invasion of the branch tip into the ECM (orange) are discontinuous in time. Scale bars, 200µm (**b**), 100µm (**c,d**), 50µm (**e**).

During branch elongation, we detected proliferative cells throughout the whole organoid (Fig. 1d) and observed the formation of filopodia-like cellular protrusions at the leading edge of the branches, hinting towards an invasive branching process (Fig 1e). This is in contrast to the non-invasive ductal elongation process driven mainly by internal cell rearrangements inside the terminal end bud, which was observed during murine mammary gland branching morphogenesis *in vivo* or in matrigel-embedded tissue fragments (18, 24). This different growth behavior could be based on the contrasting mechanical properties of the matrices. While matrigel is a purely linear material with nearly no plasticity, collagen I is highly non-linear with an initial stress softening followed by a stiffening under cyclic strain, such mechanical hysteresis loops are commonly referred as Mullins-softening (25) (Fig. 1f, Fig. S2). In our single-cell based assay we do not observe organoid growth in pure matrigel. While in mixes of collagen and matrigel the formation of only spherical clusters can be observed, highly branched structures are formed only in pure collagen gels (Fig. 1b).

As observed by means of embedded tracer particles, the elongation of each branch induced a significant long range deformation field within the ECM directed towards the branch (Fig. 1g, Fig. S1b). After 14 days of culture these local deformations added up to millimeters, which led to a macroscopic shrinkage of the collagen gel to about half its diameter. In the near-field the observed strains exhibited local heterogeneities and were highly anisotropic: deformations were the strongest directly at the tip of the branch in extension to the direction of elongation and steadily decreased with increasing angle from the branch tip (Fig. 1h, Fig. S1c, Video 1). Moreover, we observed a periodic displacement of the embedded tracer particles towards the branch with periods in the hour scale, as best seen by computing the cumulative displacement in the ECM in the near field of the branch over time (Fig. 1i, blue line, Fig. S1c, d). Analogous to the dynamic deformation field, branch elongation occurred discontinuously in time, with a back-and-forth movement of the leading cells (Fig 1i, orange line). These observations led us to hypothesize that the endogenous contractility displayed by basal, also known as myoepithelial cells, results in the detected ECM deformations, which are required for branch elongation. Since branched organoids are exclusively generated by basal, but not luminal cells (23), we wondered whether myoepithelial properties are required for the initiation of branching morphogenesis.

## Collective contractility is necessary for the formation of TDLU-like structures

Indeed, during the organoid establishment phase, we observed that small clusters consisting of just a few basal cells were already able to induce considerable deformation of the surrounding ECM (Fig. 2a). In contrast, non-contractile luminal mammary epithelial cells grew as multicellular spheres driven by a proliferative pressure without a resolvable ECM deformation (Fig. 2b). Occasionally, only short-range and localized elastic deformations appeared. To test directly whether endogenous contractility of basal cells is required for branching initiation, the Rho kinase (ROCK)-inhibitor Y-27632 was added to the culture medium from day 1 on. Indeed, branch formation was inhibited, leading mainly to the emergence of unstructured and dense cell clusters (Fig. 2c, Fig. S3a). Moreover, the inhibition of cellular contractility during the branch elongation phase on day 10 prevented further branch elongation as well as the global anisotropic contraction of the ECM as observed before. Whereas the formation of filopodia-like-protrusions was only observed in leading cells in control organoids, ROCK-inhibitor treated organoids displayed cellular protrusions in stalk cells all along the branch axis (Fig. 2d, Video 2). This observation correlated with deformations of just a few microns perpendicular to the branch in contrast to the large directed deformations observed in control organoids that emanated from the leading front in direction of branch elongation. Thus, we concluded that branch elongation is driven by the endogenous contractility of the myoepithelial cell sheet which specifically induces large anisotropic deformation fields in the ECM in front of the elongating branch. Moreover, our results suggest that branch elongation is the result of coordinated and collective myoepithelial contraction, also evidenced in the generation of filopodia-like protrusions specifically at the tip of the branch.

**Fig. 2:**
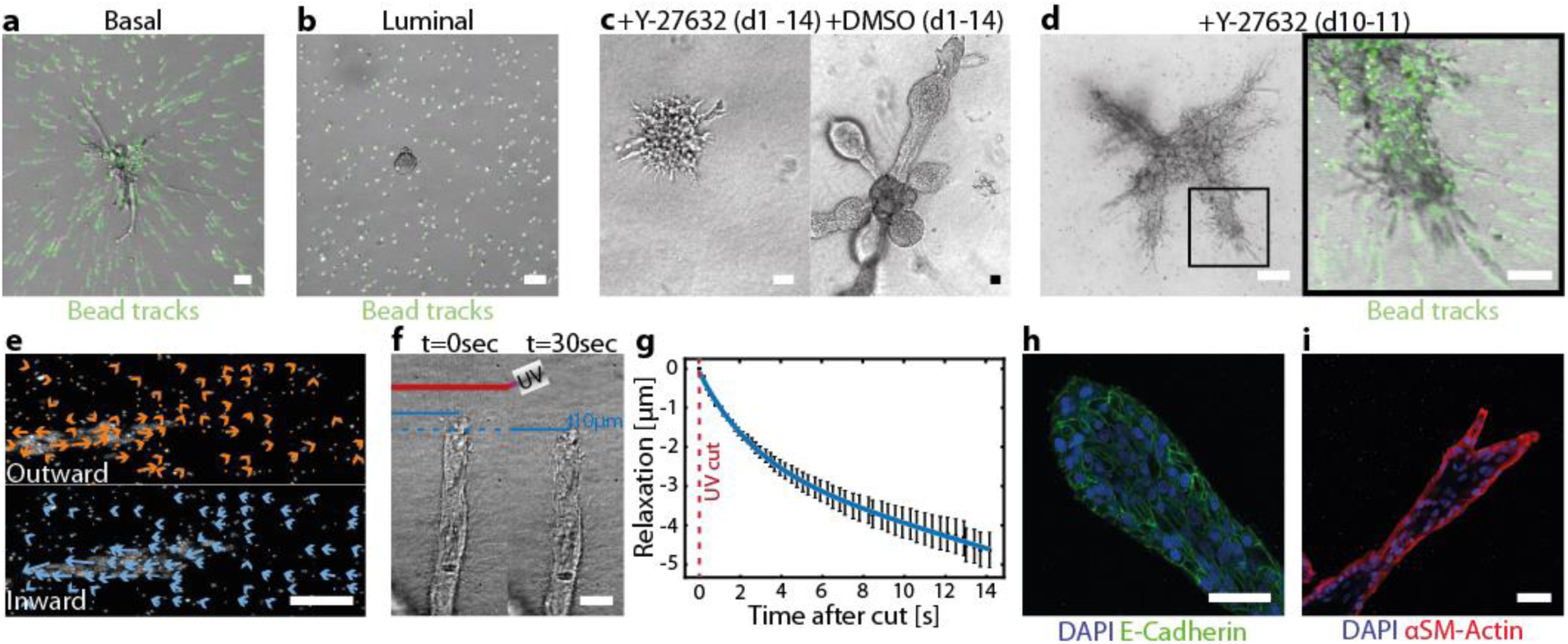
Collective contractility is necessary for the formation of TDLU-like structures. **a,b** Representative deformation field of (**a**) basal cells at day 7 over a time course of 45 hrs and (**b**) luminal cells over a time period of 19 hrs. **c** Left: Representative morphology of organoids after continuous treatment with Y-27632 (10 µM) from day 1 to 14. Right: DMSO control. **d** Left: Characteristic morphology of organoids after onetime treatment with Y-27632 during their branch elongation phase at day 10. Right: Alternated deformation field after treatment with Y-27632. n=11 organoids. **e** Internal cellular dynamics are displayed in the deformation field with cells either collectively moving outward or inward. **f** UV-cuts of the ECM in front of growing branches and following contraction of the branches towards the organoid body. n=150 branches. **g** Tracking the tip of a branch after the cut reveals a fast contraction towards the organoid body. **h,i** Representative immunostaining of (**h**) e-cadherin and (**i**) alpha smooth muscle actin (αSMA). Nuclei were labelled with DAPI. Scale bars, 50µm (**a**-**c,h,i,inset in d**), 100µm (**d**) and 15µm (**f**).

To better understand whether collagen deformation and branch elongation are mechanistically linked, we next observed cellular movements by nuclear labelling. Stalk cells moved in a collective manner inside the branch with speeds of up to 6 µm/hr, but frequently changed their direction or paused their movement (Video 1). The tip of the branch was led by a few cells, which formed filopodia-like protrusions and actively invaded the collagen network (Fig. 1e). Interestingly, the leading cells and the stalk cells occasionally switched places (Video 3), contrary to tip cells observed during endothelial sprouting (26) or tube elongation of the *Drosophila tracheal* (14). Strikingly, the direction of the net movement of the stalk cells correlated with the deformation field within the ECM in front of the branch tips. When collective cell motion was outward pointing, just small deformations in the ECM were observe. However, when stalk cells moved away from the tip towards the organoid center or showed no migration, large deformations within the ECM appeared (Fig. 2e). By contrast, no directed cell movements and deformations were observed perpendicular to the long axis of the branches. Based on these results, we speculated that the tension required for the resulting deformations we observed was applied by the whole branch and not the leading cells alone. To test this, UV laser ablation of the collagen matrix in front of invading branches was conducted (Fig. 2f). Such ablation was followed by an instantaneous relaxation of the whole branch towards the organoid body (Fig. 2g, Video 4). Accordingly, the ECM was relaxing in the opposite direction of the initial deformation field.

Together these results suggests that it is the collective nature of the cell movements within the stalk of the branch directed away from the tip which results in the observed ECM deformations in front of the branches. Thus, the tension is built up by the whole branch and is counterbalanced by the surrounding ECM. In accordance, we detected strong coupling of the cells within the branches *via* E-Cadherin (Fig. 2h). To prove the necessity of collective tension built-up, we lowered cell-cell adhesion by addition of a function-blocking anti-E-cadherin antibody (HECD1 clone) throughout the organoid culture. As a result, just spindly branched structures evolved that completely lacked alveologenesis (Fig. S3b).

## Collagen is plastically remodeled by the invading epithelium

*Via* additional immunofluorescence, overarching smooth muscle actin cables were detected in elongating branches, highlighting again the collective tension built-up (Fig. 2i). Indeed, disrupting the actin network by addition of Cytochalasin D led to a loss of tension, resulting in an instantaneous relaxation of the branch by an outward motion due to elastic counterforces of the surrounding ECM (Video 5). Yet, only a small fraction of the total deformation which accumulated during the growth process was released, which demonstrates that the observed deformations in the collagen network are predominantly plastic in nature (Fig. 3a, Fig. S4a, b). In support of these data, we observed highly aligned collagen fibers in front of the invasive branches with decreasing alignment of the fibers with increasing angle to the branch axis (Fig. 3b), which correlates to the observed deformation field. Importantly, the orientational order of the collagen network was kept in this alignment upon Cytochalasin D treatment (Fig. 3c). Finally, we observed that the collagen network far away from organoids was fully isotropic with randomly oriented collagen fibers, hinting that the fiber alignment is established by the expanding branches of the organoid (Fig. S4c). Live-cell imaging revealed that fiber alignment was induced by the collective mechanical tension produced by the cells within the elongating branches (Video 6). Thus, the fiber alignment was a direct consequence of the observed contractile deformation field and captured its history. Importantly, the tension was generated by the whole branch and not by the tip cells alone. The contractile forces of an individual basal cell do not suffice to explain the observed long ranging deformations (Fig. S4d). Moreover, tip cells did not continuously attach to one attachment site, but frequently changed their attachment sites on the collagen fibers (Fig. 3e). Yet, due to the previous growth induced alignment of the collagen fibers in front of the branch through the collective contraction and cell movement within the entire branch, orientational guidance along the axis of the branch elongation was effectively achieved.

**Fig. 3:**
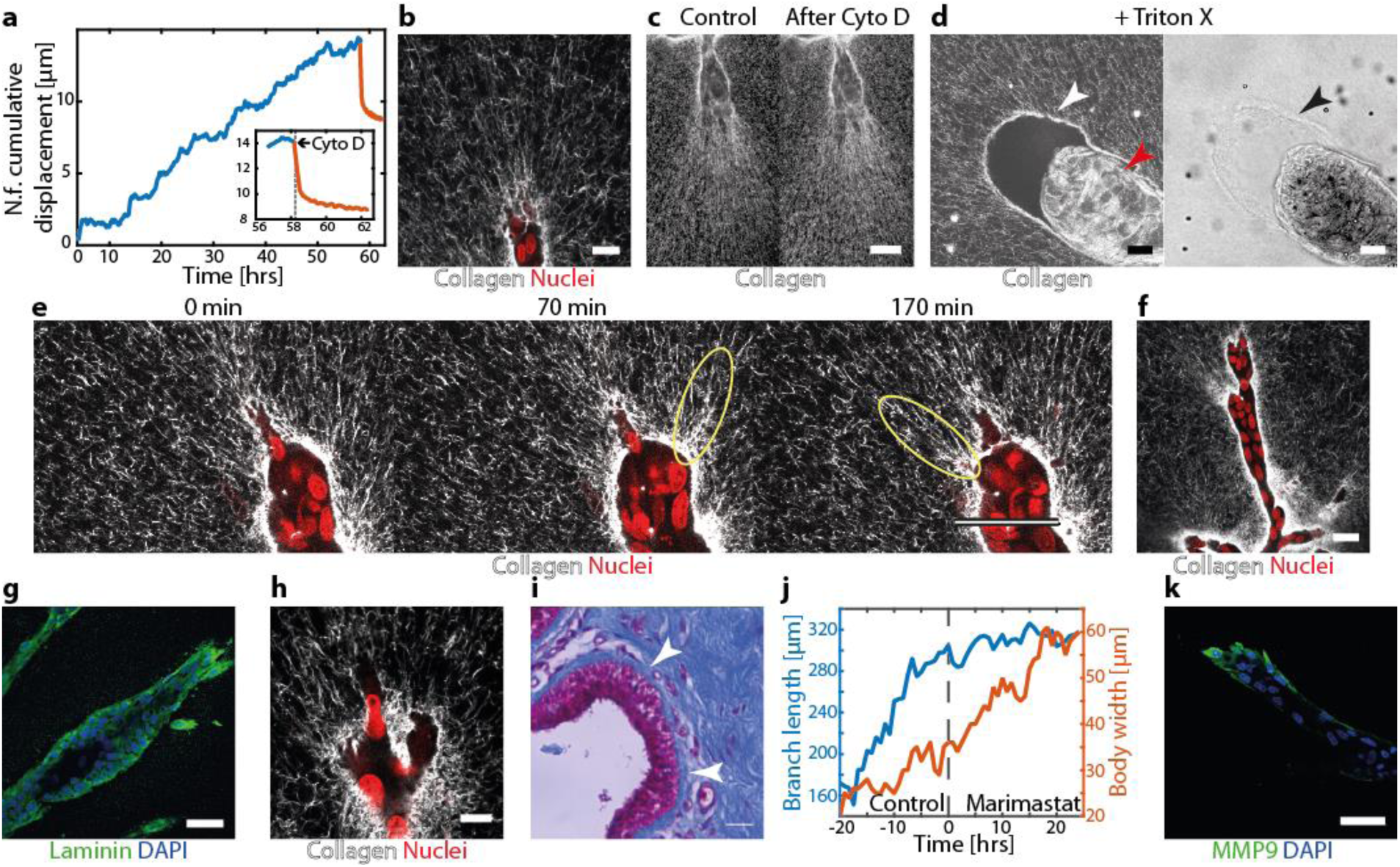
Collagen is plastically remodeled by the invading epithelium. **a** Representative behavior of cumulative displacement during the growth (blue) and relaxation upon treatment with Cytochalasin D (1 µg/ml, orange) in front of a branch. n=31 organoids. **b** Collagen fibers show alignment in front of the branches. n=5 donors. **c** Fiber alignment is conserved after treatment with Cytochalasin D. n=5 donors. **d** Organoids treated with Triton X. Left: The collagen cage (white arrow) retains its structure even after the collapse of the branch (red arrow). Right: The cage is visible in the bright field image (black arrow). **e** Time course of alternating fiber alignment of the invading branch (yellow circles). **f** Collagen accumulation around a growing branch visualized by fluorescent collagen. **g** Immunostaining of laminin showed localized expression at the cell-ECM interface. **h** During matrix invasion leading cells squeeze through the collagen network as seen by deformation of their nucleus. **i** Trichrome staining of breast tissue sections show an accumulation of collagen around the ducts (white arrows). **j** By inhibiting MMPS, branch elongation is inhibited (blue), while the body width is still increasing (orange). **k** Immunostaining of MMP9 shows localized expression in the leading cells. Scale bars: 50 µm (**b, e, g, k**), 20 µm (**c, d**), 25 µm (**f**), 10 µm (**h**).

In addition, we observed that the collagen fiber alignment resulted in an enrichment of collagen along the branch axis (Fig. 3f). This mechanically induced accumulation ultimately led to the formation of a continuous collagen cage. High-resolution microscopy revealed that this collagen cage had a porous structure with holes of an approximate size of 1 up to 3 µm (Video 7). The thickness around the organoid body had a size of around 10 µm and thinned out towards the tip of the branch with larger holes at the invasion site. Washing out the epithelial cell layer by addition of Triton X left an empty collagen cage behind (Fig. 3d). Thus, the plastic deformation of the collagen generated a collagen cage which encased the organoid and, once formed, remained mechanically stable even in the absence of cells. In addition to accumulated collagen around the epithelial branches, we detected an enrichment of cell-secreted laminin, a major component of the basement membrane (Fig. 3g). The mechanical stability of the collagen cage guides the further elongation process of branch. While at the side the dense cage prevents a further outgrowth (27), the emerging cage in the front is porous and weak enough for individual cells to squeeze through (Fig. 3h). Similar to our *in vitro* observations, a trichrome staining of human tissue cross sections showed that epithelial ducts in the human mammary gland are surrounded by Collagen I (Fig. 3i). Here, collagen is a major component of the basement membrane, which separates the mammary gland from stromal fibroblasts and is thought to act as mechanical hindrance to prevent branch outgrowth (28).

In order to test the interplay of plastic deformation and local degradation of the matrix we inhibited the activity of Metalloproteinases MMPs by the addition of Marimastat. At the beginning of the establishment phase, addition of 10µM Marimastat led to the formation of very short and thin branches (Fig. S3c). The inhibition of MMP activity in the branch elongation phase induced an arrest of invasion by the branch tip cells into the collagen, whereas formation of filopodia was still observed (Fig. 3j, blue line, Video 8). Yet, the branches were still able to build up significant tension, which in turn led to large strains in the collagen and further plastic deformation of the surrounding matrix. However, the inhibition of the MMPs hampered the extension of the branch while the cells continued to proliferate. Due to the reduced ability of branch extension continuous cell proliferation resulted in an increased cell density and a concomitant slowing of cell motility and thickening of the structures (Fig. 3j, orange line). Immunofluorescence against MMP9, a metalloproteinases previously described to have a dominant role in matrix remodeling in the mammary gland (29), revealed a localized expression at the tip of the invading branches (Fig. 3k). Based on these results, we concluded that the formation of the collagen cage resulted from a combination of mechanically induced accumulation of collagen in front of the branch and single-cell invasion of the tip cells (Video 9).

## Conclusions

Together we can summarize our presented results of the invasive branching morphogenesis into a mechanical feedback model (Fig. 4): The collective cell migration of the epithelial cells within branches leads to a tension generation. Thereby, the applied traction force of individual cells sum up *via* cell-cell adhesions, comparable to the tension build-up inside cell sheets on 2D substrates (12). The resulting tension is transmitted *via* the leading cells to the surrounding collagen. Consequently, the elastic counter force of the matrix leads to plastic deformations of the collagen such as fiber alignment in front of the branch. The invasion of the leading cells themselves relies on the local collagen degradation by MMPs, comparable to single cell migration in dense matrices (30, 31). This invasive migration leads to the built-up of a mechanical encasing of the complete branch by a stable collagen cage, which in turn facilitates the collective cell migration within the branch (Fig. 4b).

**Fig. 4:**
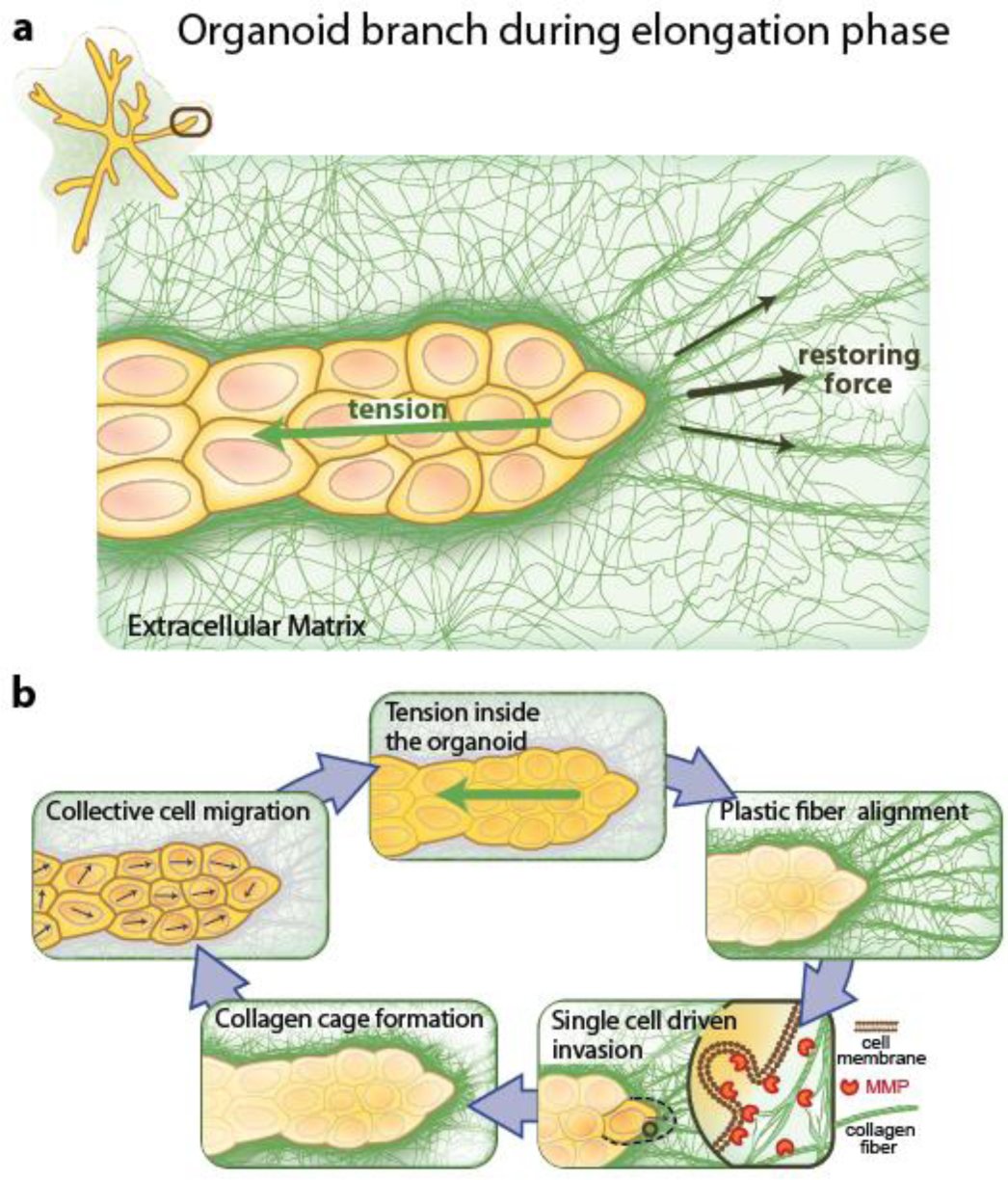
Mechanistic model for invasive branching morphogenesis. **a** There is a balance between the cellular tension of the branch and the tensile restoring forces of the ECM. **b** Cells inside the branch migrate collectively on the collagen cage and apply a traction force on it. These traction forces sum up to a tension inside the organoid, which is transferred to the ECM. As a result the collagen fibers get aligned and apply a counter force. MMPs enable the tip cells to invade the matrix, which leads to the formation of the collagen cage. Ultimately, the cells use this self-built collagen cage to migrate on.

Our results demonstrate how invasive branching morphogenesis of human mammary gland organoids is governed by the external mechanical plasticity of the ECM. Due to the universal role of branching in organogenesis, the identified mechanical feedback model paves the way to further address mechanical self-organizing principles in morphogenesis.

## Supporting information

Video 1 - Organoid growth

Video 2 - Y-27632 treatment

Video 3 - Tip cell exchange

Video 4 - Laser ablation

Video 5 - Cytochalasin D treatment

Video 6 - Collagen invasion

Video 7 - High resolution imaging of the collagen cage

Video 8 - Marimastat treatment

Video 9 - Cage formation

## Author contributions

A.R.B, C.S, B.B. and L.K.M conceived the experiment. B.B. conducted live-cell imaging and analyzed the deformation fields. L.K.M. performed immunofluorescence and drug screening. P.F. carried out the shear rheology measurements. M.K.R. applied the laser ablation experiments. F.P.H. analyzed the structure of the collagen networks. All authors contributed to the interpretation of the data and to the writing of the manuscript.

## Correspondence and request for materials

should be addressed to A.R.B and C.S.

## Materials and Methods

### Flow cytometry and fluorescence-activated cell sorting (FACS)

Single-cell suspensions of primary HMECs were stained with the following antibodies: CD31-PB, CD45-V450, CD49f-PE, EpCAM-FITC and CD10-APC (table S1-S3). 7AAD was added to the suspension for dead cell exclusion. Luminal progenitors (LP) and CD10+ basal cells (B+) were sorted as previously described (23) (Fig. S5). A subsequent re-analysis was performed to ensure the purity of the sort. FlowJo V10 software was used for post-analysis.

### Organoid preparation

Organoids were prepared as described previously (23). In short, freshly isolated human mammary gland epithelial cells from healthy women undergoing reduction mammoplasty were embedded in collagen gels (collagen type I from rat tail, Cornings) with a final collagen concentration of 1.3 mg/ml. For specific experiments pure collagen was mixed with labelled collagen in a ratio of 20:1. From day 1 to 5 cells were cultivated in mammary epithelial growth medium (PromoCell MECGM) enriched with 3 µM Y-27632 (Biomol), 10 µM Forskolin (Biomol) and 0.5% FBS. From day 7 to day 14 the media was changed to MECGM mixed with 10 µM Forskolin. Organoids were prepared from 5 different donors (Table S4).

### Collagen labelling

Collagen was fluorescently labelled with Atto 488 (Merck) according to a protocol based on a previously published protocol (32). Therefore, collagen was dialysed at 4°C to reach pH 7. Subsequent, collagen was conjugated with Atto 488 by incubating it overnight at 4°C. Further dialyse was performed for 8 hrs to remove non-bound dye, followed by an additional dialyse overnight using acid to prevent unwanted polymerization. Finally, it was stored at 4°C.

### Live-cell imaging

Live-cell imaging was done using a Leica SP8 lightning confocal microscope equipped with an ibidi gas incubation system for CO2 and O2. Organoids were labeled with sirDNA (10µM, Spirochrome AG) three hours before measurement. Organoid age and observation time was set according to the planned experiment, time between each image was kept at 10 min. For nuclei visualization samples were excited with 633 nm and detected at an emission maximum around 674 nm. Collagen fibres were either visualized by illuminating the samples with 488 nm and detecting the reflected signal or by imaging fluorescently labelled collagen. High-resolution images of the collagen cage were taken by using an implemented adaptive deconvolution algorithm.

### Data analysis

Images were processed using Fiji and Matlab. Drift correction was performed using a self-written Matlab code based on maximizing the correlation value between consecutive images by shifting images stepwise in x-y-z direction. For bead detection images were first masked by an intensity threshold. Subsequent bead position is defined by taking the center of an interpolated intensity grid. Beads touching the boundary of the image and beads below a minimum distance to each other are removed to prevent wrong trajectories. Finally, tracks were calculated by matching coordinates in three consecutive images. To calculate the cumulative displacement the individual displacement of a bead in the near field was summed up and afterwards the mean was calculated by averaging of beads in the same area. The internal flow field was calculated *via* optical flow using the in Matlab implemented Farneback algorithm.

### Immunofluorescence

Collagen gels were fixed with 4% paraformaldehyde for 15 min. For immunofluorescence, cells were permeabilized with 0.2% Triton X-100 and blocked with 10% goat or donkey serum in 0.1% BSA. Samples were incubated with primary and secondary antibodies diluted in 0.1%BSA. Antibodies and dyes used for stainings are listed in table 1, 2 and 3. DAPI was used to visualize the cell nuclei.

### Laser ablation

Laser ablation of collagen was performed with a custom-made nano-dissection setup based on the one described previously (33). Our setup includes a spinning-disc unit (CSU-X1, Yokogawa) equipped with an Andor NEO sCmos camera, a 638 nm and a 488 nm laser (Cobolt) for fluorescence confocal imaging, as well as a HAMAMATSU Orca Flash 4 for bright field imaging along a separate optical path. Cuts were done by shining a pulsed UV-laser with emission of 355 nm, 400 psec pulse duration, 72 kW peak power and 25 mW average power through a PL APO 40x 1.1 NA water objective (Leica microsystems) with a working distance of 0.65 mm. Movies were taken at frame rates between 120 fps and 200 fps. For analysis movies were processed with Fiji. The tip cell was tracked manually to measure the relaxation after the cut.

---

**Fig S1:**
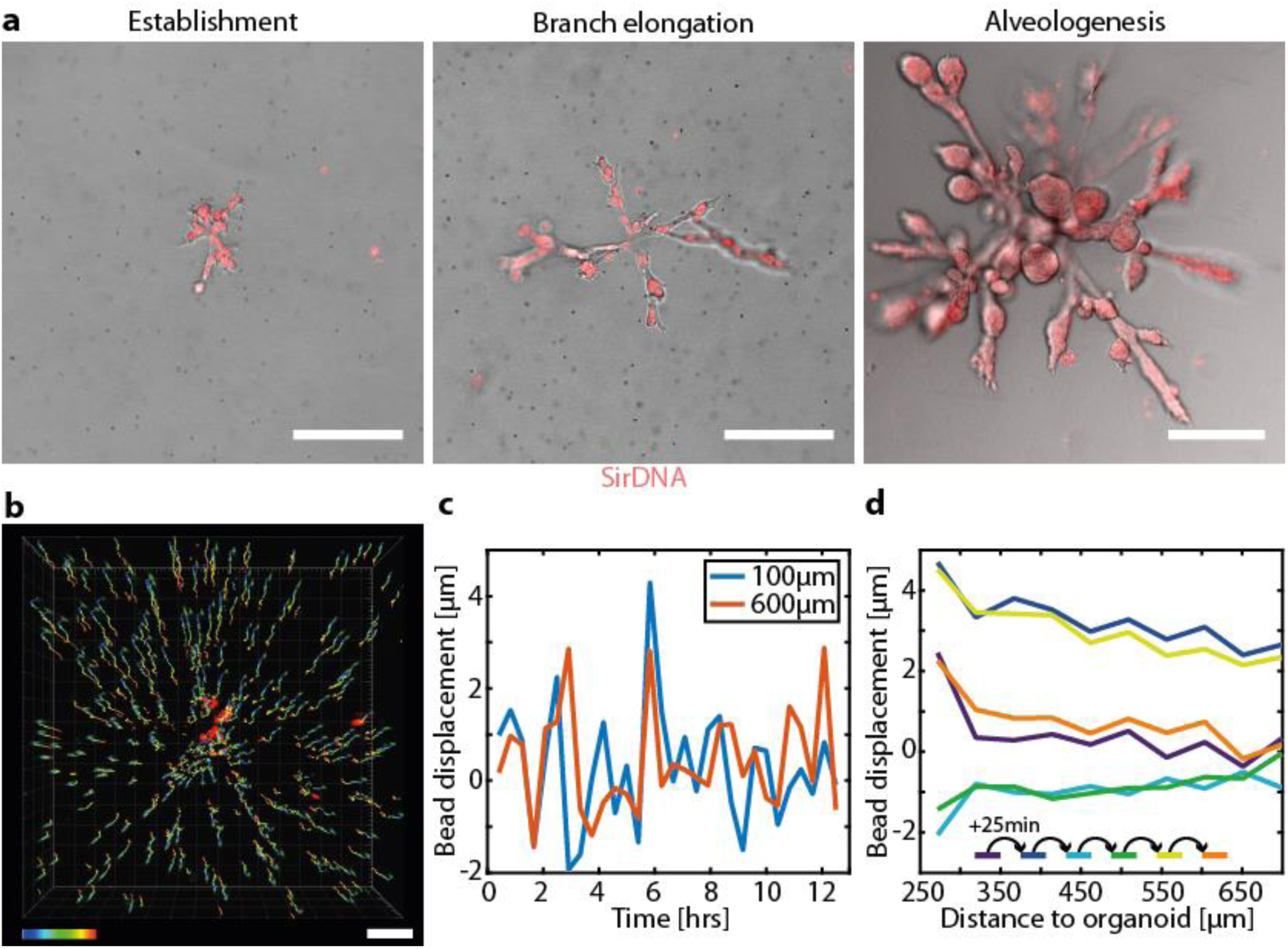
**a** Characteristic organoid morphology during different developmental stages. **b** Bead trajectories tracked over a time course of 48 hrs during the establishment phase. **c** The average displacement in the near field (100µm, blue) and far field (600µm, orange) of the branch tip shows an oscillatory behavior. **d** The bead displacement is gradually decreasing with increasing distance to the branch tip and alternates between contractions and relaxations. Scale bars: 200µm (**a**), 100µm (**b**).

**Fig. S2:**
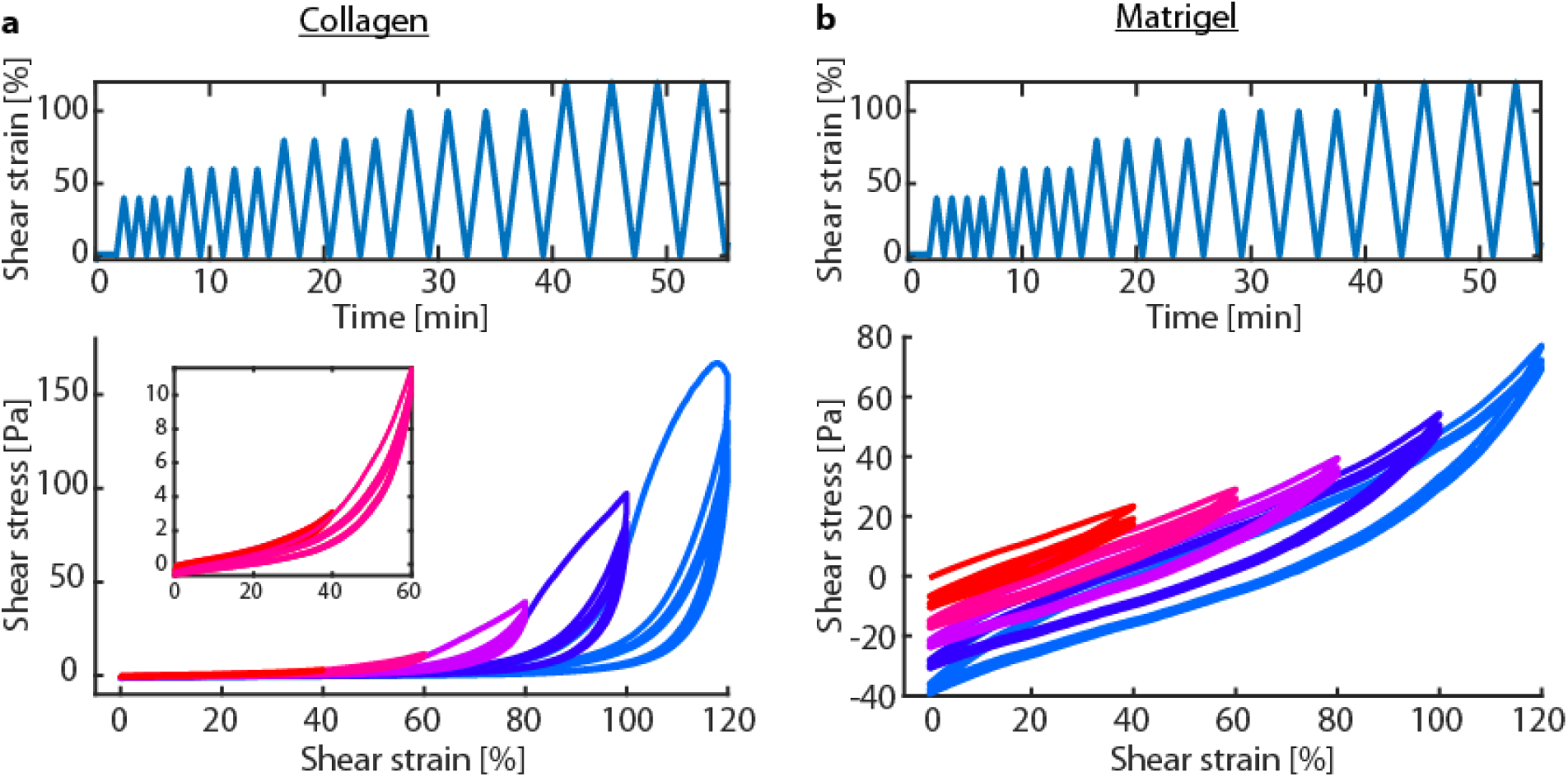
Shear rheology of Collagen and Matrigel. **a** Collagen networks show non-linear response upon shear stress. During cyclic shearing the Mullins-softening can be observed. **b** Matrigel networks show no stiffening and only little plasticity.

**Fig S3:**
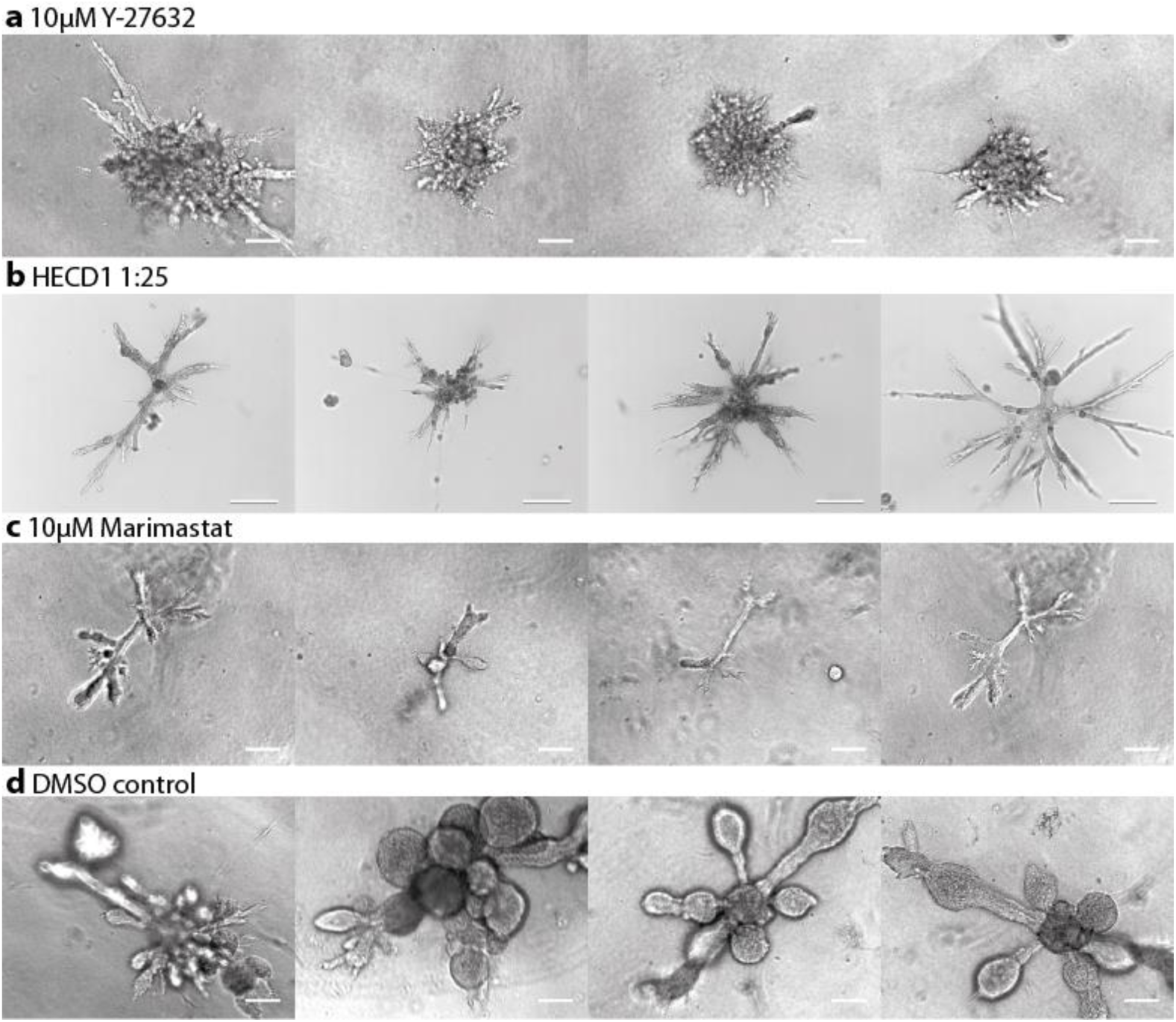
Organoid morphology after continuous drug treatment with beginning of day 7. **a** 10µM Y-27632, **b** HECD1 at a ratio of 1:25, **c** 10µM Marimastat, and **d** DMSO control. Scale bars: 100µm (**a,b,d**), 200µm (**c**).

**Fig S4:**
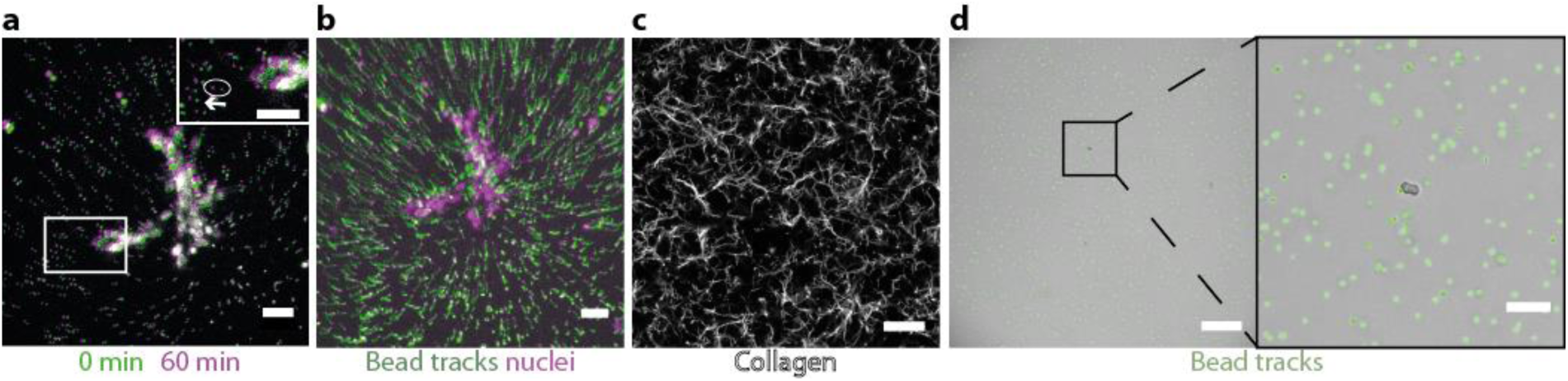
**a** Upon treatment with Cytochalasin D branches relax. Beads retract in the opposite direction of the initial deformation (white arrow). **b** The projection of the initial bead displacement prior to the treatment with Cytochalasin D shows the pronounced displacement field. **c** Far away from an organoid the collagen shows no distinct orientational order. **d** Deformation field for single basal cells for 21 hrs. Scale bars: 50µm (**a**-**c, inset d**), 200µm (**d**).

**Fig S5:**
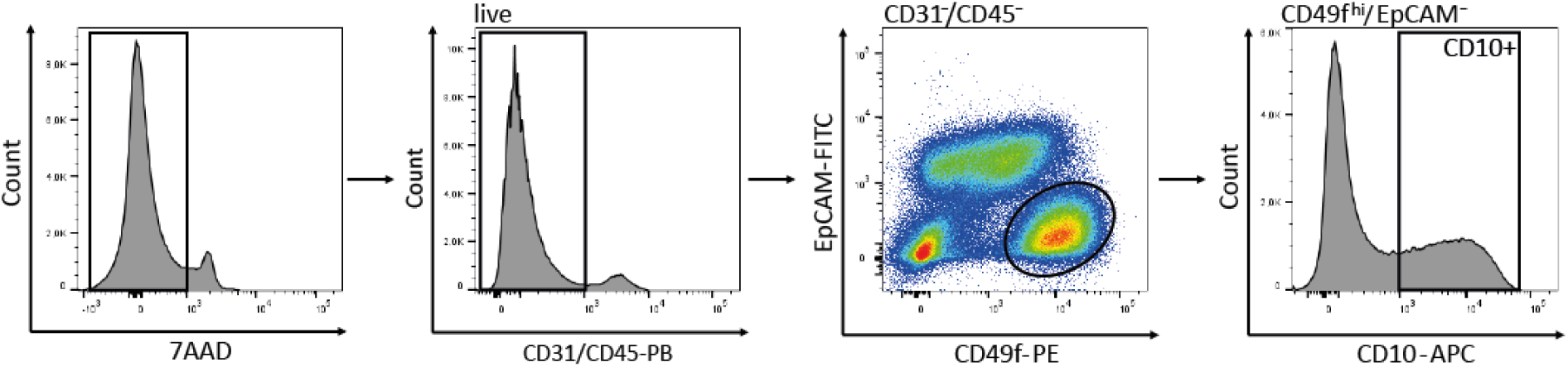
FACS gating strategy for isolation of basal mammary epithelial cells from fresh isolated human mammary epithelial cells. Dead cells (7AAD^-^ = live), hematopoietic (CD45^+^), and endothelial cells (CD31^+^) were excluded. By using the markers EpCAM, CD49f and CD10, the basal population (EpCAM^-^/CD49f^hi^/CD10^+^) was isolated.

**Video 1: Organoid growth** During the branch elongation phase beginning from day 7 the typical deformation field can be observed by tracking fluorescent beads embedded inside the collagen gel. Nuclei labelling reveals highly dynamic cell motion during the organoid growth.

**Video 2: Y-27632 treatment** Organoid behavior upon treatment with 10µM Y-27632 at day 10 shows the formation of filopodia-like structures.

**Video 3: Tip cell exchange** Stalk cells occasionally replace leading cells.

**Video 4: Laser ablation** Relaxation of a branch after laser ablation of the collagen in close proximity to the tip.

**Video 5: Cytochalasin D treatment** Relaxation field upon treatment with 10µM Cytochalasin D shows counter balance between tensile forces of the branch and restoring force of the collagen.

**Video 6: Collagen invasion** Fluorescent collagen reveals that leading cells do not continuously attach to the same collagen fibers but change their attachment sites over time.

**Video 7: High-resolution imaging of the collagen cage** 3D visualization of the collagen cage around an organoid using a deconvolution algorithm.

**Video 8: Marimastat treatment** Organoid growth upon treatment with 10µM Marimastat shows stop of branch elongation and thickening of the branches.

**Video 9: Cage formation** Collagen gets pushed to the sides by the tip cell, leading to the accumulation of collagen at the sides.

**Table S1:** Primary antibodies.

| Epitope [Clone] | Conjugation | Host | Catalog number | Supplier |
| --- | --- | --- | --- | --- |
| alpha smooth muscle actin | - | Rabbit | ab5964 | Abcam, Cambridge, UK |
| E-cadherin [HECD1] | - | Mouse | ab1416 | Abcam, Cambridge, UK |
| Gata3 [L50-823] | - | Mouse | CM405 | Biocare Medical, Concord, US |
| Ki67 | - | Rabbit | ab15580 | Abcam, Cambridge, UK |
| Laminin | - | Rabbit | L9393 | Sigma, Steinheim, Germany |
| P63 [EPR5701] | - | Rabbit | ab124762 | Abcam, Cambridge, UK |
| Phalloidin | Atto 647 | - | 65906 | Sigma, Steinheim, Germany |
| MMP9 [56-2A4] | - | Mouse | ab58803 | Abcam, Cambridge, UK |
| Integrin α6 [GOH3] | - | Rat | sc-19622 | Santa Cruz, Dallas, US |

**Table S2:**
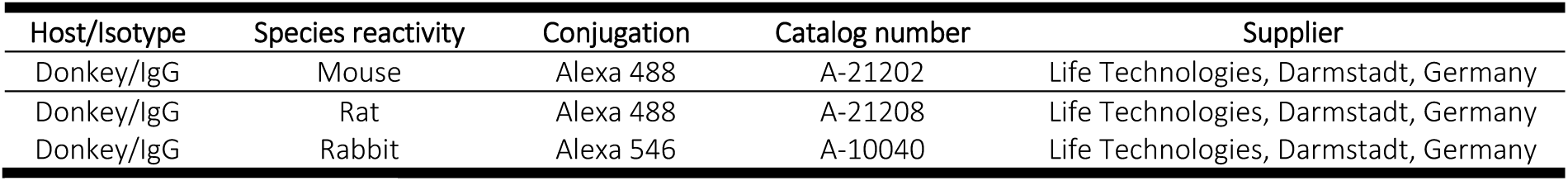
Secondary antibodies.

| Host/Isotype | Species reactivity | Conjugation | Catalog number | Supplier |
| --- | --- | --- | --- | --- |
| Donkey/IgG | Mouse | Alexa 488 | A-21202 | Life Technologies, Darmstadt, Germany |
| Donkey/IgG | Rat | Alexa 488 | A-21208 | Life Technologies, Darmstadt, Germany |
| Donkey/IgG | Rabbit | Alexa 546 | A-10040 | Life Technologies, Darmstadt, Germany |

**Table S3:**
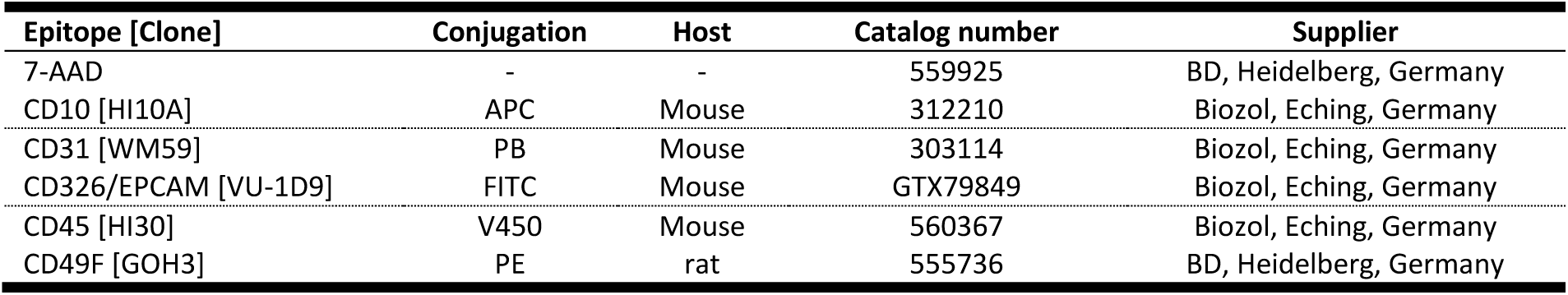
Antibodies used for flow cytometry and fluorescence activated cell sorting.

| Epitope [Clone] | Conjugation | Host | Catalog number | Supplier |
| --- | --- | --- | --- | --- |
| 7-AAD | - | - | 559925 | BD, Heidelberg, Germany |
| CD10 [HI10A] | APC | Mouse | 312210 | Biozol, Eching, Germany |
| CD31 [WM59] | PB | Mouse | 303114 | Biozol, Eching, Germany |
| CD326/EPCAM [VU-1D9] | FITC | Mouse | GTX79849 | Biozol, Eching, Germany |
| CD45 [HI30] | V450 | Mouse | 560367 | Biozol, Eching, Germany |
| CD49F [GOH3] | PE | rat | 555736 | BD, Heidelberg, Germany |

**Table S4:**
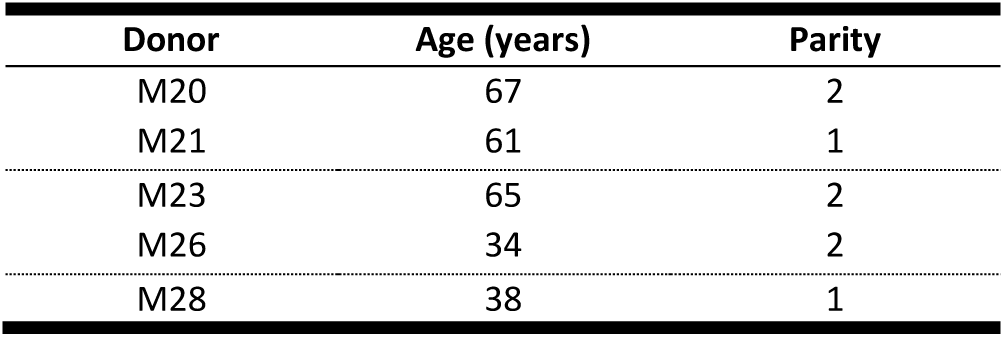
Reduction mammoplasty donors.

| Donor | Age (years) | Parity |
| --- | --- | --- |
| M20 | 67 | 2 |
| M21 | 61 | 1 |
| M23 | 65 | 2 |
| M26 | 34 | 2 |
| M28 | 38 | 1 |

